# A role of territory formation in disruption of heterologous pairings in *Drosophila* male meiosis

**DOI:** 10.1101/2020.08.28.272658

**Authors:** Christopher A. Hylton, John E. Tomkiel Dean

## Abstract

Pairings between heterologous chromosomes in meiosis can lead to nondisjunction and the production of aneuploid gametes. To minimize these aberrant outcomes, organisms have evolved mechanisms to disrupt such improper pairings prior to orientation and segregation. In the male fruit fly, *Drosophila melanogaster*, bivalents segregate to distinct nuclear domains in prophase I, and it has been proposed that the formation of these distinct territories may play a role in disrupting interactions between limited homologies on heterologous chromosomes. To test this, we used fluorescent in situ hybridization to examine pairing between the X chromosome and *Dp(1;3)* chromosomes in which a segment of the X had been transposed to chromosome 3. We found that 120kb of homology was sufficient to insure nearly complete pairing but was not sufficient to direct merotelic segregation of the paired elements, suggesting that such pairings were being disrupted. We compared the perdurance of X / *Dp(1;3)* pairings to that of X / *Dp(1;Y)* pairings (in which homologs are paired),and found that heterologous pairings were disrupted at a higher frequency at the S2b stage of prophase I, the stage at which territory formation is initiated. Our results support the model that movement of bivalents into distinct domains in prophase I provides a mechanism to disrupt pairings between limited regions of homology, and thus may be one means of preventing improper segregation of heterologs in this organism.

## INTRODUCTION

Meiosis consists of two cell divisions following a single round of replication that result in the production of four haploid daughter cells from a single diploid parent cell. Errors in meiosis produce aneuploid gametes, and subsequent zygotic gene dosage imbalances can lead to inviability or severe genetic syndromes. A number of events must occur with high fidelity to ensure error-free meiotic divisions. The first, or reductional, meiotic division requires that homologous chromosomes find their partners, pair and conjoin, orient and attach to opposite poles, then coordinately separate to precisely half the number of chromosomes in each daughter cell. Partner recognition and pairing have long been recognized as critical steps in the reductional division, yet the mechanisms by which these events occur remain elusive. The criteria that determine how homologous sequences recognize each other and join together, and how distinctions are made between similar sequences on homologs versus heterologs are also largely unknown.

The male Drosophila is a useful organism to study pairing, as meiosis occurs without crossing over, and in the absence of the complexity of the regulating and establishing exchanges, arguably provides a simplified model. The fly has only four chromosome pairs, which can readily be visualized in male meiosis. In addition, the sex chromosomes and fourth chromosomes can be distinguished from the two major autosomes by size and morphology. These properties have allowed for features of meiotic chromosome behavior to be defined by direct examination of condensed chromosomes. Such observations have been informative in revealing aspects of meiotic pairing and conjunction: in particular, differences between sex chromosomes and autosomes.

The X and Y pair and conjoin at a specific locus, the tandem rDNA repeats embedded in heterochromatin located near the centromere of the X and at the base of the short arm of the Y (Cooper 1959; Ritossa 1976; McKee and Karpen 1990). Each of these clusters contain 200-220 copies of the rDNA cistrons (Ritossa 1976), and these repeats constitute the majority of sequence homology between the X and Y. While it has not been ruled out that the entire rDNA cistrons contribute to pairing, they are not required for it. Transgene studies have shown that a repeated 240 bp intergenic spacer (IGS) region within the promoter of the 18s rDNA genes is sufficient for both pairing and conjunction (McKee, Habera, and Vrana 1992; McKee and Karpen 1990). Interestingly, while an rDNA transgene on the X can efficiently pair with and segregate from the Y chromosome, an autosomal insertion of the same transgene cannot (McKee and Karpen 1990). This difference may suggest that there is something special about the sex chromosome pairing sites that limits their function in a sex-chromosome-specific manner.

For the autosomes, euchromatic, but not heterochromatic, sequence homologies have been shown to be sufficient for mediating pairing and conjunction (Yamamoto 1979; Hilliker, Holm, and Appels 1982). Compound autosomes sharing euchromatic homology are frequently associated at prometaphase I and metaphase I, whereas those sharing only heterochromatic homology are not. Although these results have led to the conclusion that heterochromatin is not involved in autosomal pairing, they allow for the possibility that pairing can occur at heterochromatin, while conjunction can only be established at euchromatic sequences. The specificity of conjunction may be determined by three proteins that may act in a complex; Stromalin in meiosis (SNM), Mod(mdg4) in meiosis (MNM) (Thomas et al. 2005) and Teflon (Tef) (Arya et al. 2006; Tomkiel, Wakimoto, and Briscoe 2001). SNM and MNM are required for all chromosome pairs, whereas Tef is autosome-specific. The binding sites for this putative complex on the autosomes have not been determined.

In contrast to the localized pairing sites of sex chromosomes, autosomal pairing and conjunction sites appear to be numerous and distributed throughout euchromatin. These conclusions were derived from the behaviors of 2-Y transpositions (*Tp(2;Y)s*) that were examined at mid to late prophase I and prometaphase I. Segments of chromosome 2 euchromatin transposed onto the Y chromosome pair and conjoin with their intact chromosome 2 partner, while a single transposition of chromosome 2 heterochromatin does not (McKee, Lumsden, and Das 1993). In general, the longer the duplicated chromosome 2 material, the more proficient the association with and segregation from the intact homolog.

This study employed relatively large segments of translocated chromosome 2 material, and the limits of the material within each transposition at the time could only be grossly defined by cytology of salivary gland chromosomes. Thus, individual pairing sites could not be mapped. Additionally, conclusions about which sequences were effective at pairing were conflated by the inability to distinguish pairing from conjunction. Nonetheless, they suggest that euchromatic sequences could mediate both pairing and conjunction.

To separate the sequence requirements for pairing versus conjunction, it is necessary to examine pairing directly in early prophase. Direct observations of associations between lacI-GFP-labeled integrated *LacO* sequences, reveal the pairing program in this organism to be more complex than previously thought. Pairing is completed in early prophase I, but somewhat surprisingly, by mid-prophase I, pairing at individual loci appears to be resolved. Chromosome-level associations are somehow maintained between homologs, as bivalents reside in discrete nuclear domains at this stage (Vazquez, Belmont, and Sedat 2002). How homologs remain associated at this stage is somewhat of a mystery, as centromeres appear to be separate (Vazquez, Belmont, and Sedat 2002) as does pericentric heterochromatin (Tsai, Yan, and McKee 2011). The sequestering of bivalents in individual domains may be important not only for maintaining connections between homologs, but also for disruption of inappropriate pairings between heterologs (Vazquez, Belmont, and Sedat 2002).

Recently, we used fluorescent *in situ* hybridization (FISH) to directly examine early prophase pairing between endogenous sequences. We found robust pairing between an rDNA-deficient *In(1)sc^4L^sc^8R^* X and *Dp(1:Y)* chromosomes, in which a segment of X euchromatin was inserted on the Y (Hylton et al. 2020). This result indicates that pairing between sex chromosomes is not limited to the rDNA IGS sequences, but rather can occur at other homologous sequences as well.

Here, we ask if X chromosome euchromatin is limited to mediating pairing between the X and the Y, or can X euchromatin also establish pairing with an autosome. We use FISH to directly assay pairing between the X and transpositions of X material onto chromosome 3 (*Dp(1;3)s*) We use genetic assays to ask if such putative pairings can establish conjunction and direct merotelic segregation. Lastly, we ask if prophase I domain formation I disrupts heterologous pairings, by examining the timing of dissolution of heterologous versus homologous pairing.

## MATERIALS AND METHODS

### *Drosophila* Stocks and Crosses

*Drosophila* were raised on a standard diet consisting of cornmeal, molasses, agar, and yeast at 23°C. All *Dp(1;3)* stocks were obtained from the Bloomington Stock Center (Gramates et al. 2017).

### Genetic Assays of Meiotic Chromosome Segregation

Segregation of a *Dp(1;3)* chromosome from an intact X was monitored by crossing *y w sn* / Y; *Dp(1;3)* / + males to *y w sn* females. *y w sn* / *y w sn; Dp(1;3)* / + females were crossed to *y w sn* / Y males to control for viability.

A segregation value S (McKee, Lumsden, and Das 1993) was calculated as the proportion of euploid progeny from *y w sn* / Y; *Dp(1;3)* / + fathers in which the duplication segregates from the X, adjusted for viability difference using segregation data from *y w sn* /*y w sn; Dp(1;3)* / + mothers. Chi-squared tests were performed to determine if S differed from 0.5, the expectation of random assortment.

S = (X + 3 from Fathers) / [(X + 3 from Fathers) + [(Y + 3 from Fathers) * [(X + 3 from Mothers) / (Y + 3 from Mothers)]]]

### Probe Design

Triple-labeled probes pools were generated to selected sequences at a density of 10 probes/Kb and a complexity of ~10,000 probes per pool (Arbor Biosciences, Ann Arbor, MI). ATTO-594 oligonucleotide probes were generated to hybridize to 1,000 Kbp present on the X and the following regions of X sequences duplicated on the *Dp(1;3)* chromosome 3s:

*Dp(1;3)RC017:* X salivary gland chromosome bands 3D5-3E1, bp 3543803-3606837. *Dp(1;3)RC029:* X salivary gland chromosome bands 12A4-12D2, bp 13824546-14001084. *Dp(1;3)RC035:* X salivary gland chromosome bands 17F2-18A2, bp 18900731-19062922.

A triple-labeled ATTO-488 probe was generated to bp 20368577-21368577 (56F-57F) on chromosome 2.

### FISH

Testes from larvae or pharate adults were dissected in Schneider’s Drosophila media (GIBCO BRL, Gaithersburg, MD). Tissue was transferred to a drop of Schneider’s on a silanized coverslip and gently squashed onto a Poly-L-Lysine coated slide (Electron Microscopy Sciences, Hatfield, PA). Coverslips were immediately removed after freezing in liquid nitrogen. Tissue was fixed in 55% methanol / 25% acetic acid for 10 min followed by 10 min dehydration in 95% ethanol. Slides were processed immediately or stored for up to 1 week at 4°C.Slides of testis tissue were processed for FISH using a slight modification of the protocol as described in Beliveau, Apostolopoulos, and Wu (2014), as reported in Hylton et al. (2020). S1-S2a (10 to 20 μm) and S2b (>20 to 30 μm) spermatocytes were selected based on size, and signals were scored as paired when within 0.8 μm as described in Beliveau, Apostolopoulos, and Wu (2014).

### Data Availability

All strains are available on request. The authors affirm that all data necessary for confirming the conclusions of the article are present within the article, figures, and tables.

## RESULTS / DISCUSSION

### *Dp(1;3)s* pair with but do not effectively segregate away from the X

To ask if X euchromatic homologies duplicated on chromosome 3 could pair, conjoin, and direct segregation from an intact X, we used a series of *Dp(1;3)* chromosomes that contain X euchromatin of varying lengths and arise from different locations on the X (Figure 1). These X transpositions were created by transformations of BACs using phiC31-mediated site-directed insertion, and thus have the advantages of both being molecularly defined and of all residing at the same position on chromosome 3L (65B2) (Venken et al. 2010). The *Dp(1;3)s* contain X homology ranging in length from 21 to 177 kb. These lengths are roughly the same size as the X euchromatin duplicated on *Dp(1;Y)s* (~120 Kb of X homology) previously found to be sufficient for pairing and segregation from the X (Hylton et al. 2020), and thus might be useful for drawing comparison to such results.

**Figure 1.**
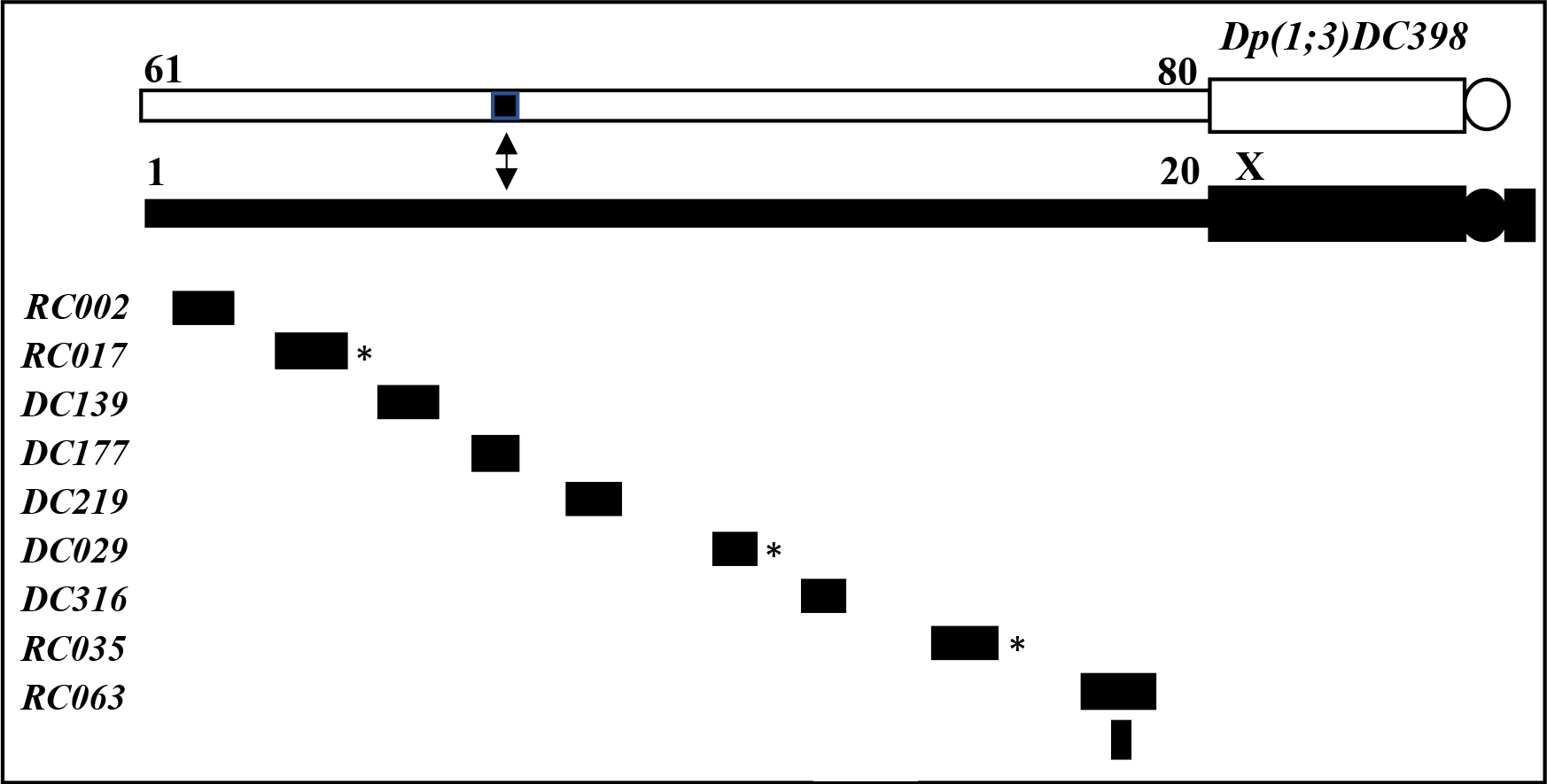
Regions of X euchromatin duplicated on each *Dp(1;3)* chromosome. The X is indicated by black boxes, 3L is indicated by open boxes. Numbers above euchromatin refer to salivary gland bands. * Pairing was examined by FISH for these duplications.

Pairing of a subset of three *Dp(1;3)s* and the intact X were monitored using FISH probes that hybridized to the X euchromatic sequences duplicated on the *Dp(1;3)*. These sequences were selected because of their various positions on the X (distal, medial and proximal) and because they were of a size that could be robustly and reproducibly detected by FISH. Pairing is reported to be complete in spermatocytes by early S1 of prophase I (Vazquez, Belmont, and Sedat 2002), and we identified cells at this stage based on size (between 10 and 20 μm (Cenci et al. 1994)) and the pairing status of a control chromosome 2 probe. The pairing assay used here is described in Material and Methods, and at length in Hylton *et al*. (2020).

Each of the three *Dp(1;3)s* analyzed paired with the intact X in greater than 90% of the cells (Figure 2, Table 1). The frequency of pairing was not affected by the location of the homology on the X. These results confirm those of our previous study, in which we found that that sequences capable of pairing are distributed throughout X euchromatin (Hylton et al. 2020), and they extend our previous analysis by showing that such pairings can occur between heterologous chromosomes. While it was known that interactions between heterologs could occur from observations of late prophase associations between chromosome 2 and T(Y;2)s (McKee, Lumsden, and Das 1993), it was not known that they occurred at such high frequencies. The efficacy of these pairings suggests that during the early prophase, there may be no mechanism that limits interactions to homologs in this organism.

**Figure 2.**
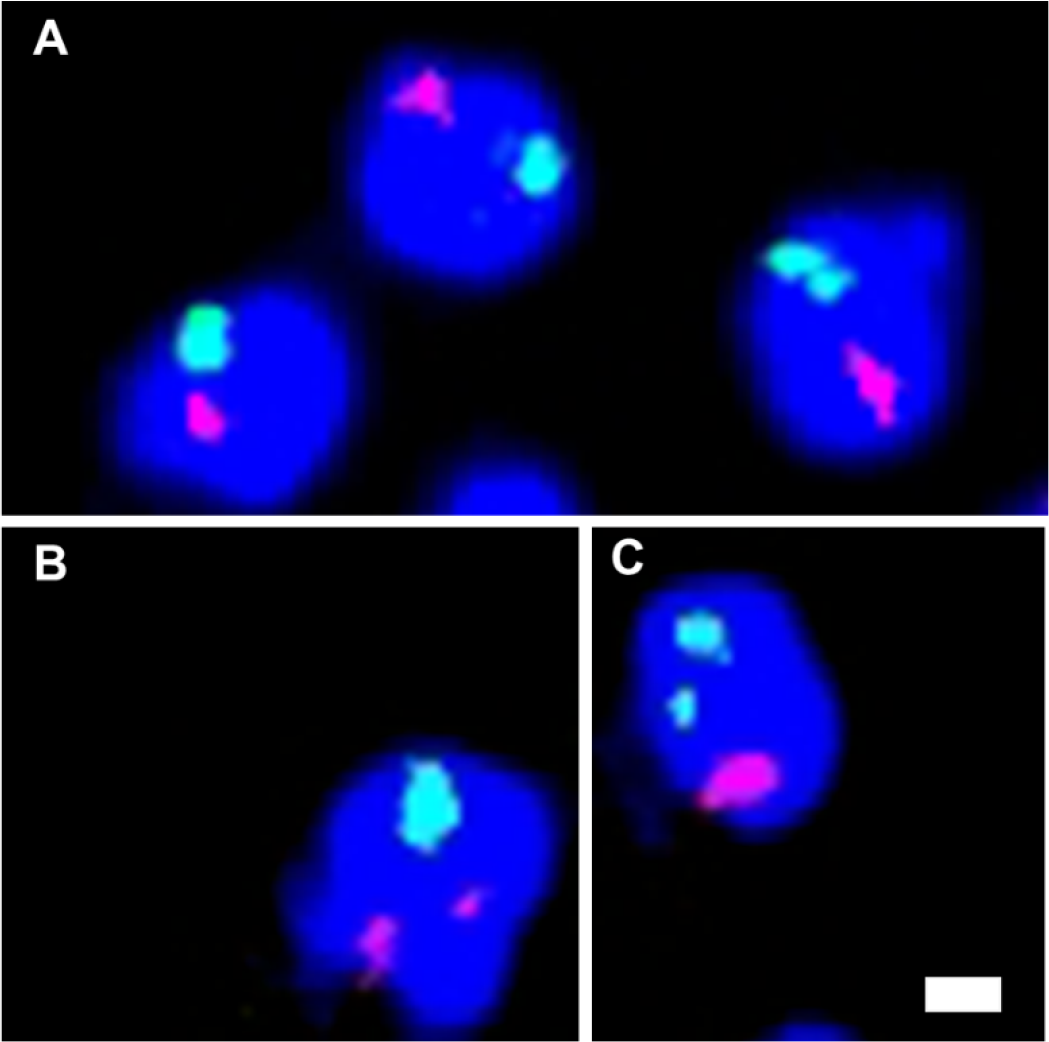
FISH examination of X / *Dp(1;3)* pairing in DAPI-stained S1-S2a primary spermatocytes. The X chromosome probe is labeled red and the chromosome 2 probe is labeled green. (A) Paired X / *Dp(1;3)* chromosomes and paired chromosome 2 bivalents. (B) Unpaired X / *Dp(1;3)* chromosomes and a paired chromosome 2 bivalent. (C) Paired X / *Dp(1;3)* bivalent and unpaired chromosome 2s. Scale bar = 2 μm.

**Table 1.** X / *Dp(1;3)* pairing in S1-S2 primary spermatocytes

| *Dp(1;3)* | X Sequences Duplicated | (kbp) | # Cells | % Paired |
| --- | --- | --- | --- | --- |
| *Dp(1;3)RC017* | 3,543,803.. 3,706,696 | (163) | 212 | 92.9 |
| *Dp(1;3)RC029* | 13,824,546.. 14,001,084 | (177) | 204 | 93.1 |
| *Dp(1;3)RC035* | 18,900,731.. 19,062,922 | (162) | 383 | 90.3 |

To determine if these pairings were effective in establishing conjunction and directing merotelic segregation of the paired elements, we performed genetic tests of segregation. As we were not limited by probe expense or signal detection for these tests, we were also able to assay the segregation of additional *Dp(1;3)* chromosomes. We selected duplications that originated from non-overlapping regions spanning the X euchromatin (Figure 1). From X / Y; *Dp(1;3)* / + fathers, we asked how often the *Dp(1;3)* chromosome segregated with the Y (i.e. to the opposite pole of the intact X). To eliminate effects of the duplication on viability, only the euploid progeny classes were considered in this analysis (i.e. those arising from X; + and Y; + sperm). To control for any other potential viability differences between the resulting classes, a segregation value S was calculated in which ratios were adjusted by those of identical progeny generated from X / X*; Dp(1;3)* / + females (See Materials and Methods). In these females, the *Dp(1;3)* must segregate with one of the two identical Xs, so any variability in the recovery of X; + sons versus daughters should reflect only viability differences.

Of the ten duplications tested, seven showed segregation from the X at frequencies that were significantly greater than random (Table 2). These included the smallest duplication, *Dp(1;3)DC398*, bearing only 21 kb of X homology. Other larger duplications (e.g. *Dp(1;3)RC002* bearing 140 kb of homology) showed no evidence of segregation from the X. This difference suggests that the distribution of sequences in X euchromatin required for conjunction is nonrandom. This notion is supported by the speckled distribution of MNM on autosomes in MI, suggesting a nonuniform distribution of conjunction complex proteins (Thomas et al. 2005).

**Table 2.**
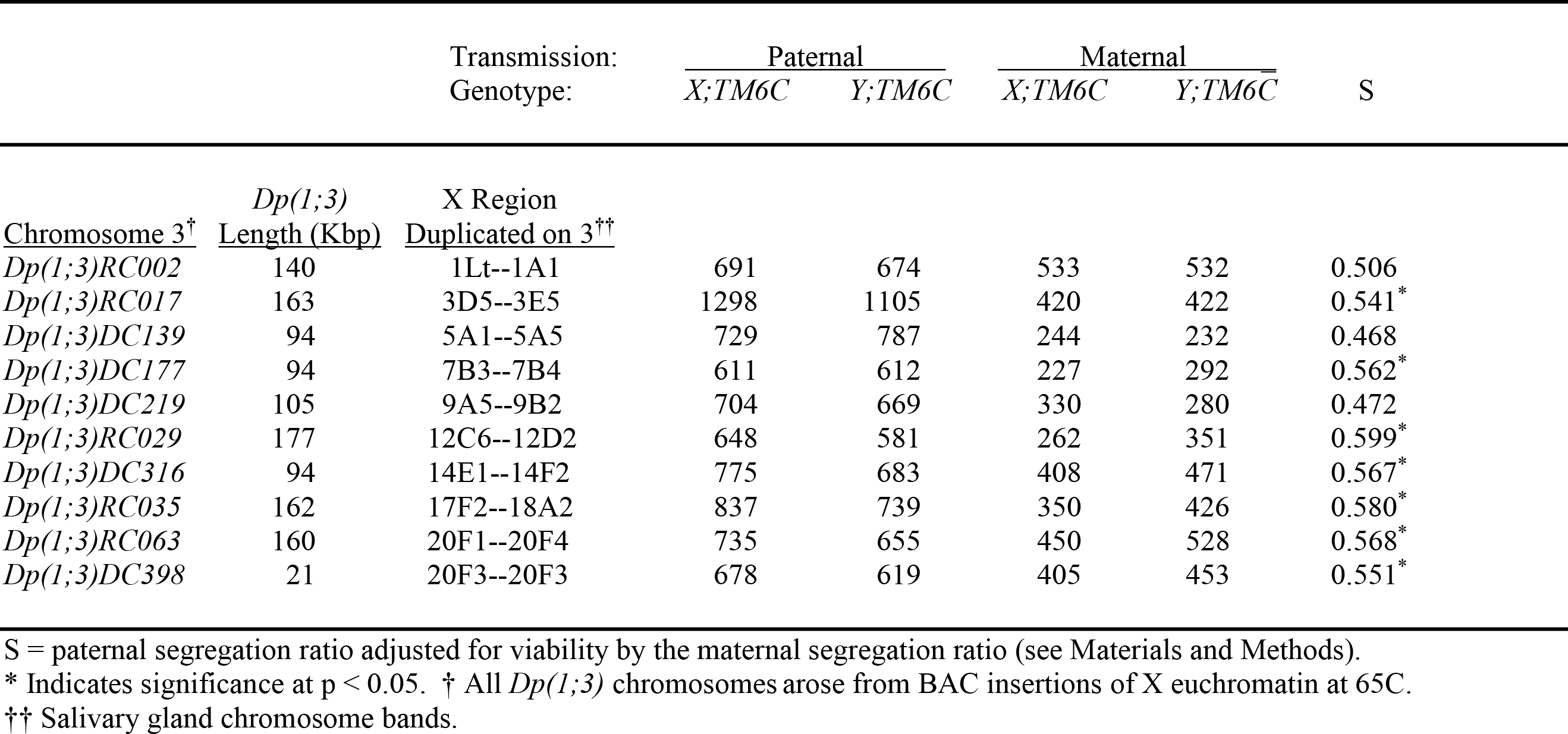
Segregation of *Dp(1;3)* chromosomes from an intact X chromosome.

Although 7 of 10 *Dp(1;3)s* segregated from the X, the frequencies of segregation in all tests were minimal, with the best being only 10% greater than random. These frequencies are similar to segregation frequencies of the *Tp(2;Y)s* from chromosome 2 (McKee, Lumsden, and Das 1993). For the three Dps for which we have pairing data as well as segregation data, it is clear that pairing was not sufficient for merotelic segregation in most meioses.

The inability of the *Dp(1;3)s* to segregate from the X is exceptionally different than the outcomes we observed for pairings between *Dp(1;Y)s* and the X. Previously, we found that *Dp(1;Y)s* bearing comparable amounts of X euchromatin segregate much more frequently from an rDNA-deficient X, in up to 90% of meioses. This variation suggests an inherent difference between the abilities of *Dp(1;3)s* and *Dp(1;Y)s* to segregate away from the X. One possible explanation for the variation is that a mechanism exists to disrupt pairings between heterologous chromosomes prior to orientation.

### *Dp(1;3)* and X pairings are resolved earlier than are *Dp(1;Y)* and X pairings

Previous work examining pairing at inserted *LacO* arrays found that homologs separate at these sites during S2b of prophase I (Vazquez, Belmont, and Sedat 2002). At this stage, the formation of three chromosome domains can first be visualized around the periphery of the cell, each of which contains a major homologous chromosome pair (Cenci et al. 1994; Vazquez, Belmont, and Sedat 2002). It has been theorized that the formation of chromosome domains sorts out heterologous pairings while maintaining proper homologous pairings (Vazquez, Belmont, and Sedat 2002). We wondered if we could determine if the disruption of the X and *Dp(1;3)* pairings is concomitant with domain formation.

To ask this, we assessed two *Dp(1;3)*s and two *Dp(1;Y)*s for their ability to remain paired with the X at S2b. We chose these rearrangements as they showed differences in segregation patterns; *Dp(1;3)RC017* and *Dp(1;3)RC035* segregated from the X in fewer than 60% of meioses (Table 2), whereas *Dp(1;Y)BSC11* segregated from the X in 70% of meioses, and *Dp(1;Y)BSC76* segregated from the X in almost 90% of meioses (Hylton et al. 2020).

A number of considerations had to be taken into account to accurately assess pairing behavior. First, as cells progress from S1 to S2b, chromosomes begin to unpair, but they do so somewhat asynchronously. That is, we expected that in some cells, unpairing of chromosome 2 might precede unpairing of the X sequences, whereas in others, the reverse may be true. Second, we also had to account for differences in pairing frequencies of different Dps examined, as we wanted to consider only cells in which we were sure pairing of the X and Dp had occurred. Third, by stage S3, pairing is lost between all sites examined (Vazquez, Belmont, and Sedat 2002), so any informative differences between heterologous and homologous pairings had to be observed before this stage was reached.

To address these issues, we only scored cells in which the X sequences were paired (Figure 3). We scored cells at both S1-S2a, when pairing was expected to be maximal, and at S2b when dissolution of pairing was expected to have initiated but before it was complete. Among these cells, we monitored the frequency at which the chromosome 2s were unpaired. Our rationale was as follows: if a particular X and Dp remained paired longer on average, then the fraction of cells with unpaired chromosome 2s would be greater. On the other hand, if the X and Dp pairing was disrupted early relative to chromosome 2, then the fraction of cells with paired X but unpaired chromosome 2s would be smaller.

**Figure 3.**
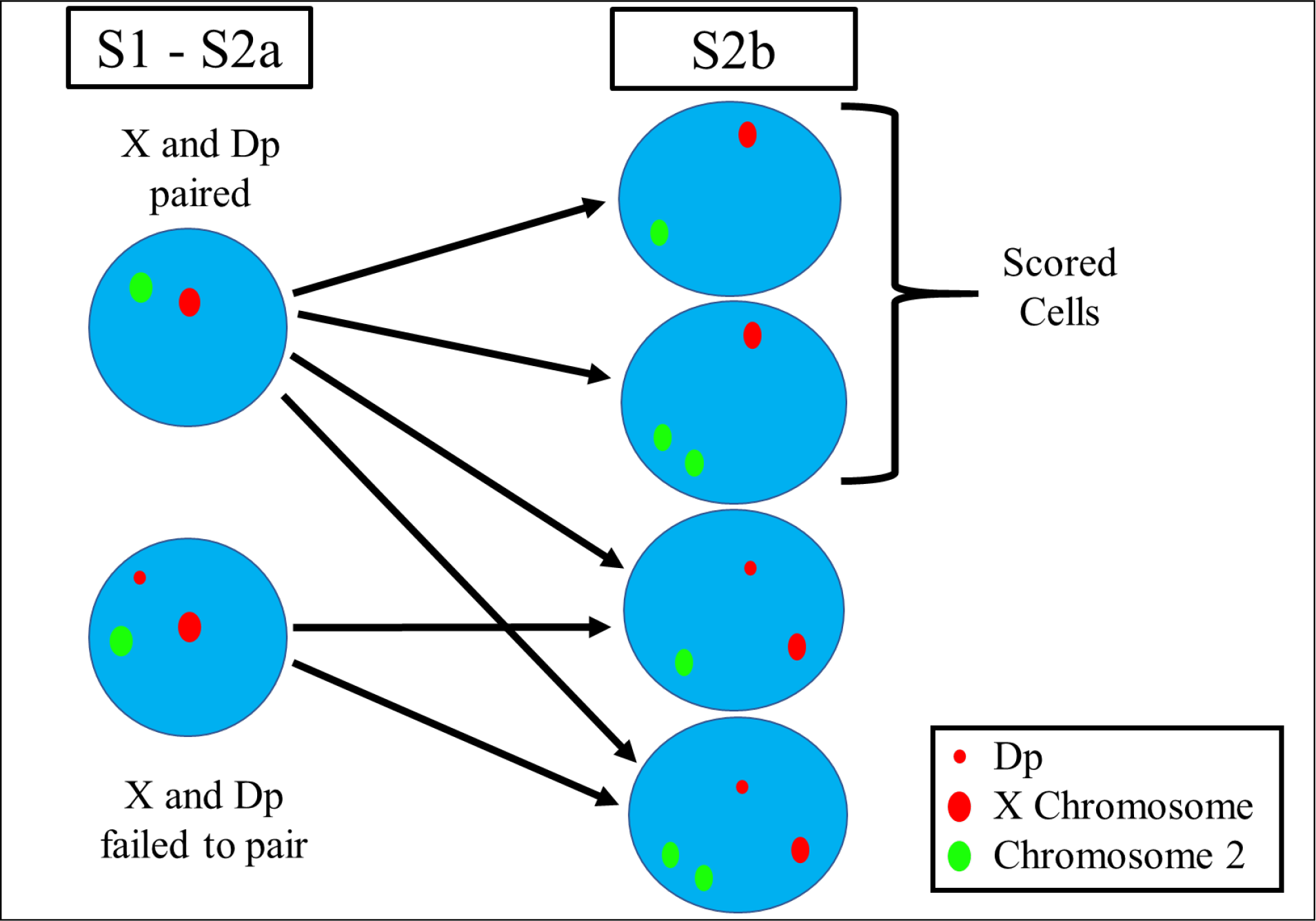
Possible outcomes at S2b. The diagram shows the predicted pairing configurations of X (red) and chromosome 2 (green) probes in S2b, based on the pairing status in S1. The X chromosome probe signal on the X is larger than on the *Dp(1;3)* because the probe binds ~1 Mbp on the X and a shorter sequence (120-300kb) on the Dp (Table 1). To limit our analysis to S2b cells in which the X probe sequences had paired in S1, FISH-probed S2b cells were scored only if the X and Dp were paired (See Results and Discussion).

In a small frequency (<10%) of cells deemed to be at S1-S2a based on their size, the X probes were paired while chromosome 2 probes were unpaired (Figure 3). It is possible that the probed autosomal regions of these cells may have never paired. Their frequency, however, was greater among X / *Dp(1;Y)* individuals than in X / *Dp(1;3)* individuals (Figure 4), and there is no prior reason to expect a difference in chromosome 2 behavior in these different genotypes. Given that there is a continuum of cell sizes at each stage, we suggest that these few cells may have already begun progression to S2b in which unpairing had been initiated. The difference in frequency between the genotypes suggests a difference in behavior of the *Dp(1;3) vs. Dp(1;Y)* chromosomes. This difference was much greater in larger S2b cells (>20 to 30 μm) (Cenci et al. 1994) where unpairing had further progressed. At this stage, there was a significantly larger fraction of cells in which the X probes were paired but the chromosome 2 probes were unpaired in X */Dp(1;Y)* than in X / *Dp(1;3)* genotypes (Figure 4). The two X / *Dp(1;3)* genotypes exhibited similar behavior, as did the two X / *Dp(1;Y)* genotypes. We interpret these results to mean that the dissociations between the *Dp(1;3)* and X chromosomes occurred early relative to dissociation of the control chromosome 2 probe, whereas dissociations between the *Dp(1;Y)* and X chromosomes occurred later.

**Figure 4.**
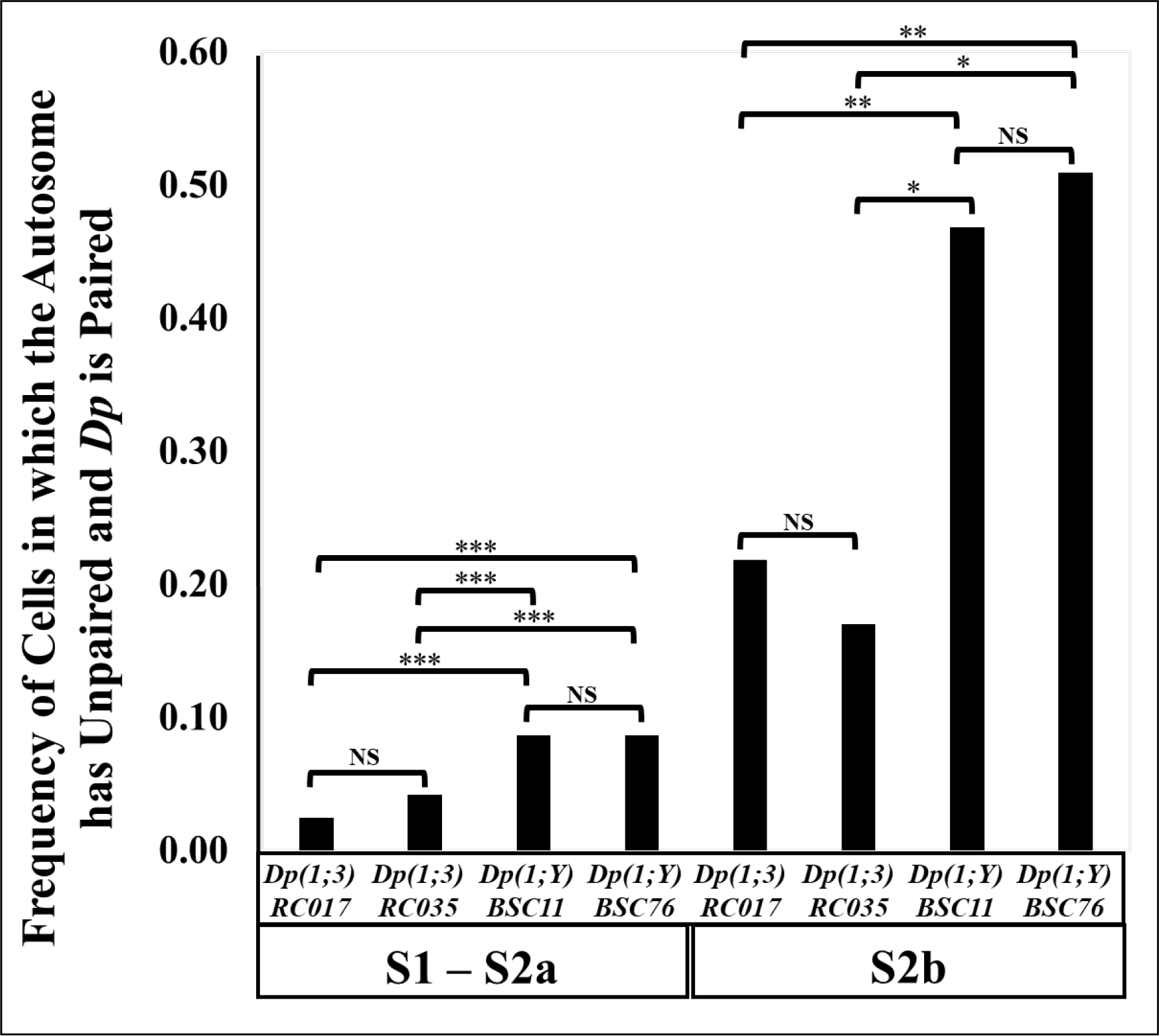
Pairing at X euchromatin on *Dp(1;Y)s* vs. *Dp(1;3)s* relative to pairing at a chromosome 2 control sequence. Frequencies of X / *Dp(1;3)* pairings and X / *Dp(1;Y)* pairings in cells where the chromosome 2 bivalent had transitioned to an unpaired state. NS = No significance. Significant difference at * p < 0.05; ** p < 0.01; *** p < 0.001.

The earlier separation of heterologs, concomitant with domain formation, might explain why pairings between the X and a *Dp(1;Y)* are better at directing merotelic segregation; they conjoin and migrate to the same domain. Perhaps, the paired duplicated X euchromatin on *Dp(1;3)* is unable to conjoin with the X or if it does conjoin, the connections are not strong enough to maintain connections when heterologs migrate to different domains.

Our results provide evidence supporting the hypothesis of Vazquez, Belmont, and Sedat (2002) who suggested that one function of domain formation may be to disrupt inappropriate pairings between heterologs by quarantining homologs to different sites in the cell. Using our FISH pairing assay, we tested a range of X euchromatin sequences, and all paired with high fidelity regardless of whether the homology was between homologs or heterologs (Hylton et al. 2020). However, there are significant differences in the abilities of these sequences to conjoin and/or segregate. It is plausible that there may be a threshold of conjunction sites required for partners to remain conjoined during domain formation, and that the *Dp(1;3)*s tested may have been below that threshold.

Little is known about how or why domain formation mechanics are initiated and how homologs remain in their own territory throughout prophase I. Since *Drosophila* male meiosis lacks many aspects of the traditional meiotic script including formation of the synaptonemal complex, recombination and resulting chiasmata between homologs, it is possible that a separate mechanism evolved to ensure accuracy of homolog pairing. Alternatively, chromosomes movements into domains may be an adaptation of a conserved underlying process. In *C. elegans*, heterologous interactions are proposed to be disrupted by chromosome movements generated by cytoskeletal connections across the nuclear membrane. A KASH/SUN-domain protein complex connects chromosomal “pairing centers” across the nuclear membrane to the cytoskeleton, and chromosome movements are thought to jostle apart inappropriate pairings between heterologs while maintaining homolog pairings (Sato et al. 2009; MacQueen and Villeneuve 2001). It remains to be seen if the movement of chromosomes in *C. elegans* and the formation of chromosome domains in *Drosophila* are based on similar molecular components. Only a single mutation, *dany*, has been described that disrupts domain formation in fly spermatocytes, and *dany* has no known relationship to components of the *C. elegans* system that generate meiotic prophase chromosome movement (Trost et al. 2016).

## CONCLUSIONS

Results obtained here suggest that all homologies are capable of meiotic pairing in *Drosophila* male meiosis, with the caveat that the duplications examined for pairing were relatively large, the smallest being ~120 kb. Examining pairing between smaller homologies by FISH may be informative, although such studies may be limited by the ability to detect shorter probes.

We found no indication that pairing was suppressed between heterologous chromosomes, with frequencies of X pairings being greater than 90% for all three *Dp(1;3)s* examined, suggesting that there is no mechanism to limit pairing to homologs.

While pairing may be promiscuous, the ability to pair was not directly related to the ability of paired chromosomes to segregate from one another. Thus, there is a difference in the ability of different regions to direct segregation, which may be based on differential binding of conjunction complex proteins.

As pairings occur very efficiently between sequences on heterologs, a mechanism for disrupting heterologous pairings is critical in this organism. We provide evidence that the variation between the ability of paired sequences to direct segregation is related to domain formation. Pairings between the X and *Dp(1;3)s* in a significant number of cells were resolved concomitant with domain formation, whereas fewer pairings were X and *Dp(1;Y)* were resolved at this stage. The migration of bivalents to separate nuclear domains may provide the forces needed to separate poorly conjoined pairings between heterologs while maintaining appropriate homolog connections.

## FUNDING

This work was supported by National Institutes of Health grant R15GM119055 to J.E.T and the University of North Carolina at Greensboro and a UNCG Graduate Student Research Award to C.A.H.

